# Lack of Reproducibility of Gulf War Illness-like Behavioral Effects in War-related Chemical Agents and Stressor Exposure Rat Models

**DOI:** 10.1101/2025.11.06.687069

**Authors:** Rashelle Lashley, Joshua Brian Foster, Xueqin Wang, Albert John Muhleman, Samuel Weaver, Edison Lin, Liching Lai, Chien-liang Glenn Lin

## Abstract

Gulf War illness (GWI) is characterized by a range of neurological and neuropsychiatric symptoms, including chronic pain, fatigue, mood problems, and cognitive issues. The pathophysiology is associated with war-related environmental chemical exposure and psychological stress, which are known to disrupt glutamatergic synaptic structure and function. The goal of this study was to evaluate the efficacy of the investigational new drug LH-001, a candidate for enhancing tripartite glutamatergic synapses, in a rat model of GWI. We began by attempting to induce GWI-like phenotypes in rats using a previously published protocol involving exposure to specific chemical agents and mild stress. However, these efforts were unsuccessful, as the expected GWI symptoms were not observed. We then performed two protocol modifications—using more potent chemical exposures and higher stress levels—but were still unable to produce the desired phenotypes. Our results highlight significant issues with the reliability of current GWI rat models. A comparative analysis of other studies suggests that GWI-like behavioral effects, when induced in rats by chemical agents and stress, show a broader lack of reproducibility across different laboratories. This calls into question the robustness of current rat models for GWI and poses challenges for preclinical drug development.

## Introduction

Gulf War illness (GWI) is a chronic, multi-symptomatic condition affecting approximately one-third of veterans—around 250,000 people—who served in the 1990–1991 Gulf War [1]. GWI presents a wide range of symptoms that include fatigue, gastrointestinal disorders, and widespread pain, among others [2]. Neurological problems, including headache, sleep difficulty, anxiety, depression, memory deficit, and difficulty in concentration, are commonly reported symptoms [3–6]. Although the exact etiology of the disease remains unclear, it is widely thought that these clinical symptoms stem from a combination of war-related stress and exposure to a range of toxic chemical agents [7–10]. These chemical agents include pyridostigmine bromide (PB, administered as a prophylactic treatment against organophosphate nerve agents); pesticides (such as permethrin (PER) and N,N-diethyl-meta-toluamide (DEET)); chemical warfare agents (such as sarin); burn pits; byproducts of oil well fires; and depleted uranium. Current treatments for the complex disorder of GWI focus on managing symptoms; however, understanding its mechanisms is critical for developing more effective therapies.

Growing evidence indicates that exposure to GW-related chemical agents can impair the structure and function of glutamatergic synapses in the central nervous system [11]. PB, pesticides, and sarin are acetylcholinesterase inhibitors [12]. Under stress conditions, these chemicals can enter the brain and inhibit acetylcholinesterase activity. This results in elevated acetylcholine levels, leading to excess glutamate release and subsequent excitotoxic damage and synaptic dysfunction [13–14]. Air pollutants from burn pits can penetrate the respiratory system, enter the bloodstream, cross the blood-brain barrier, and enter the brain. Subsequently, they can induce neuroinflammation, oxidative stress damage, excitotoxicity, and disruption of glutamatergic synapses [15–16]. Depleted uranium can enter the brain and cause altered glutamate uptake and glutamate receptor expression, neurochemical imbalances, oxidative stress, and neuroinflammation [17]. Exposure to Agent Orange has been linked to several neurological problems including cognitive and neuropsychiatric problems, primarily through the dioxin contaminant 2,3,7,8-tetrachlorodibenzo-p-dioxin which can damage glutamatergic synapses and alter glutamate transmission [18–19]. Moreover, mounting evidence indicates that stress can cause dysregulation of glutamate transmission and losses of dendrites and their synapses [20–21]. Taken together, military-related toxic exposures can impair glutamatergic synapses, resulting in long-term structural and functional alterations in the brain and leading to neurological problems.

LDN/OSU-215111 is a novel compound that can enhance the structure and function of tripartite glutamatergic synapses—a functional unit composed of a presynaptic terminal, a postsynaptic terminal, and an astrocytic process. We previously tested LDN/OSU-215111 in a mouse model of GWI [22]. In this model, mice were exposed to three chemical agents associated with GW, including PB (1.3 mg/kg, orally), PER (0.13 mg/kg, dermally), and DEET (40 mg/kg, dermally). Following chemical administration, mice were exposed to two different unpredictable stressors, a procedure designed to replicate the multiple, unpredictable psychological and environmental stress factors faced by GW veterans in a war zone. In exposed mice, we observed anxiety-like behaviors, cognitive impairments, and muscle weakness. LDN/OSU-215111 treatment reversed these behavioral and muscular phenotypes. It also normalized associated hippocampal changes, including elevated glutamate, impaired glutamatergic synapses, interneuron loss, decreased neurogenesis, and altered excitatory and inhibitory transmission [22].

In this study, we extended our previous findings in mice by validating the compound’s beneficial effects in rats. We began with the procedure from Parihar et al. [23], which used the same GW chemical cocktail (1.3 mg/kg PB, 0.13 mg/kg PER, and 40 mg/kg DEET) but with a milder, 5-minute restraint stress (hereinafter referred to as “low dose/low stress” protocol). Disappointingly, this exposure did not produce any reported GWI-like phenotypes. A subsequent experiment was performed with a significantly increased dose of chemical agents (13 mg/kg PB; 2.6 mg/kg PER; and 215 mg/kg DEET) combined with 5-minute restraint stress (hereinafter referred to as “high dose/low stress” protocol) [24–25], which also failed to produce the expected GWI-like phenotypes. Afterwards, we used the same exposure protocol as our previously used mouse model [22]; it consisted of 1.3 mg/kg PB, 0.13 mg/kg PER, and 40 mg/kg DEET, followed by two unpredictable stressors (hereinafter referred to as “low dose/high stress” protocol). Unfortunately, we observed neither the behavioral phenotypes associated with GWI nor the significant pathological changes previously reported. Results of this study suggest that GWI-like effects are not reliably reproducible in rats. Potential factors that contribute to the discrepancy between other reports and our study are discussed.

## Materials and Methods

### Animals

Male Sprague Dawley rats (Charles River) were used in all cohorts. Rat starting weights ranged from 225-275 g, at an estimated age of 7-8 weeks. Rats were left to habituate in the vivarium for one week before beginning the stress and chemical exposure protocol. Vivarium conditions were kept at a steady temperature (approximately 70°F) with a 12-hour room light/dark cycle. Rats were housed in groups or pairs depending on size and provided ad libitum access to food and water, except on days that required food or water deprivation. All procedures were done between 8 am and 6 pm. All procedures were approved by the Institutional Animal Care and Use Committee of The Ohio State University following the National Institutes of Health Guide for the Care and Use of Laboratory Animals.

### Chemical Agents

Chemical agents used for GWI induction in rats included the following: PB (Sigma-Aldrich Cat# P9797) dissolved in distilled water, PER (Sigma-Aldrich Cat# 45614) dissolved first in Dimethyl Sulfoxide (DMSO; Sigma-Aldrich Cat# 276855) and then diluted in 70% ethanol (diluted from 200 proof ethanol, Fisher scientific Cat# BP28184), and DEET (Sigma-Aldrich Cat# D100951) diluted in 70% ethanol. Batches of chemical agents were made fresh weekly and stored at 4°C when not in use. PER solution was sonicated for at least five minutes before use to minimize precipitation. All vehicles and chemicals were warmed to room temperature before use.

### GWI Induction Protocols

The induction protocols used in this study are shown in **Table 1**. Rats were weighed at the start of each week. The average body weight within groups was determined to be used for preparing the chemical agents. All rats were treated with chemical agents or vehicle 5 days a week for 4-5 weeks. GWI rats were treated via oral gavage with either a low (1.3 mg/kg) or high (13 mg/kg) dose of PB (500 µl), followed by topical treatment on a shaved region of the neck with either a low (0.13 mg/kg) or high (2.6 mg/kg) dose of PER (200 µl), and either a low (40 mg/kg) or high (215 mg/kg) dose of DEET (200 µl). After chemical exposure, rats were left to rest in their home cage for five minutes. They were then exposed to five minutes of restraint stress in plastic DecapiCones (FisherSci, Cat. # NC9745006) or two different unpredictable stressors (**Table 2**). Control rats were treated via oral gavage with 500 µl of distilled water, followed by two topical applications of 70% ethanol (200 µl each, 400 µl total) on a shaved region of the neck. Control rats were not subjected to stress. After exposure, rats rested in their home cages for 4–5 weeks. During this time, rats were handled only once weekly for body weight measurements and cage changes. Following the resting period, rats were treated with either a vehicle (saline) or the compound LH-001. Behavior testing began four or six weeks later (8–10 weeks post chemical and stress exposure). The compound treatment continued until euthanasia.

**Table 1.** GWI induction protocols.

| Protocol | Chemical and stress exposure<br>(every weekday for 4 or 5 weeks) |  |  |  | Vehicle or<br>LH-001<br>(time to receive<br>post-exposure) | Behavior testing<br>(time to conduct<br>post-exposure) |
| --- | --- | --- | --- | --- | --- | --- |
|  | PB<br>(mg/kg) | PER<br>(mg/kg) | DEET<br>(mg/kg) | Stress |  |  |
| **low dose/**<br>**low stress** | 1.3 | 0.13 | 40 | 5-minute restraint<br>stress (5 weeks) | 4-5 weeks | 8-9 weeks |
| **high dose/**<br>**low stress** | 13 | 2.6 | 215 | 5-minute restraint<br>stress (5 weeks) | 4-5 weeks | 8-9 weeks |
| **low dose/**<br>**high stress** | 1.3 | 0.13 | 40 | 2 unpredictable<br>stressors (4 weeks) | 4 weeks | 10 weeks |

PB: pyridostigmine bromide; PER: permethrin; DEET: N,N-Diethyl-meta-toluamide

**Table 2.** Unpredictable stressors.

| Stressor | Description of stressor |
| --- | --- |
| **Heat stress** | Rats were placed in cages warmed to 37°C via a heat lamp and left there for 1 hour. Cage temperatures were monitored with a thermometer. Rats were checked every 15 minutes. |
| **Cold stress** | Rats were placed in barren cages sitting on ice (temperature ranges between 8-14°C) for 1 hour. Rats were checked every 15 minutes. |
| **Food deprivation** | Food access was removed from home cage for 24 hours. |
| **Water deprivation** | Water access was removed from home cage for 24 hours. |
| **Single housing** | Rats were housed in novel cages, alone, and without enrichment for 24 hours. |
| **Light cycle disruption** | Rats in home cages were illuminated with a lamp for the duration of one full 12-hour dark cycle. |
| **Wet cage at a 30° tilt** | Novel, barren cages containing a layer of water (the greatest depth is ~0.5 cm) were secured at 30° tilt. Rats were placed in these cages for 4 hours. |
| **Restraint stress** | Rats were placed in well-ventilated clear plastic tubes measuring 2.5 cm in diameter and 25.5 cm in length with caps on either end. Rear caps contained a 1 cm x 1 cm hole for tail accessibility. Rats were restrained for 1 hour. |
| **Forced swim** | Rats were placed in a 6-gallon cylindrical fish tank with ~30 cm of 23-25°C water. Rats were left to swim for 10 minutes and monitored for the duration of the stressor and several minutes after. Rats were dried and returned to their home cage. |
| **Stroboscopic lighting** | Rats were subjected to randomized stroboscopic lighting in a dark room for 1 hour. Rats were checked every 15 minutes. |
| **Wet cage** | Rats were placed in novel, barren cages that had a layer of water ~3 cm deep for 3 hours. Rats were checked every 1 hour. |

### Compound and Treatment

LH-001, which was previously named LDN/OSU-0215111-M3 [26] and is a metabolite of LDN/OSU-0215111 [22], was used in the current study. It is an advanced compound being developed for clinical use. LH-001 (>95% purity) was highly soluble in water. After the resting period, rats were treated with either saline vehicle or LH-001 (10 mg/kg) every other day via oral gavage for 4 weeks. All animals continued to receive treatment in this manner until euthanasia.

### Behavior Assessments

Rats were subjected to a series of behavior tests to assess anxiety and cognitive functions. All behavior testing was performed between 10 a.m. and 6 p.m. Prior to each experiment, animals were habituated to the testing room ambient conditions for at least 30 minutes. The rats were then single caged for 10 minutes immediately before behavioral testing. After testing, each animal was returned to its home cage. The apparatus was cleaned with 70% ethanol between each animal to disrupt scent cues, and all apparatuses were cleaned with 10% bleach at the end of each day. Data were captured and analyzed using SMART software (Version 3.0.05, Panlab Harvard Apparatus®).

### Open Field

Rats were placed in the center of a 100 cm x 100 cm x 40 cm open field apparatus. Open field trials ran for 10 minutes under low light (approximately 30 Lux) and were recorded. The percentage of time spent in the center and total distance traveled were measured and analyzed.

### Dark and light

Dark and light testing was done using a 100 cm x 100 cm x 40 cm open field apparatus. A divider and lid were placed to create a 100 cm x 70 cm light field and a 100 cm x 30 cm dark field within the apparatus. The divider contained a small hole for rats to pass between the two fields. The light field was set to approximately 350 Lux. Rats were placed in the light field, away from the entrance to the dark field, and allowed to explore for 10 minutes. The percentage of time in the light field and latency time to the dark field were measured and analyzed.

### Elevated plus maze

This apparatus consisted of a plus shaped platform measuring 122 cm x 122 cm and stood 1 m above the ground. Two arms were closed while two arms were open. General lighting was set with the open arms being brighter (approximately 80 Lux) than the closed arms (approximately 30 Lux). Rats were placed on the end of an open arm and allowed to explore for 10 minutes. The percentage of time spent in the open arms was measured and analyzed.

### Novelty-suppressed feeding

Novelty-suppressed feeding was done using Kellogg’s brand Froot Loops cereal. Rats were introduced to the cereal in their home cages two days prior to testing. Rats were food deprived overnight right before the test trial. These tests were done in a 100 cm x 100 cm x 40 cm open field under bright light conditions, approximately 80-90 Lux. A small dish of cereal was placed in the center of the field. Rats were allowed to freely explore the open field and interact with the cereal for 10 minutes. At the end of the 10 minutes, rats were returned to their home cages, and the dish of remaining food was weighed. The latency to first interaction with the cereal and the total percentage of cereal eaten were measured and analyzed.

### Novel object location

Rats were first habituated to the empty 100 cm x 100 cm x 40 cm arena for 10 minutes per day over two days. On day three, two identical objects were placed into the upper right and upper left regions of the arena. The rats were given 10 minutes to freely interact with the objects. On day four, the upper right object was moved to the lower right of the arena. Rats were again given 10 minutes to interact with the objects. Time spent with each object was measured. A discrimination index [(novel % time – familiar % time) / (novel % time + familiar % time)] was calculated to determine the relative interaction preference. Positive values indicate more interaction with the novel location object, negative values indicate more relative time spent interacting with the unmoved object, and zero represents equal interaction with both objects. Memory deficits are indicated by a decreased discrimination index relative to controls.

### Novel object recognition

This test was an extension of the novel object location test. After the familiar object was moved on day four, the unmoved familiar object was replaced with a similar, novel object on day five. Animals were again allowed to freely explore the two objects in the arena for 10 minutes. Memory was then analyzed by generating a discrimination index, as described above, for the novel versus familiar object, with cognitive deficits defined as discrimination index scores significantly lower than controls.

### Barnes maze

The Barnes Maze apparatus was a 48-inch diameter round table (∼1 m above the ground) with twenty 8 cm diameter holes equidistantly cut into the perimeter. One of these holes was chosen to have an escape box affixed below it that the animals were trained to find to escape the bright, open arena. Eight visual cues were placed around the table. Lighting was set to 80-90 Lux to induce stress. During the training phase, rats were placed in the center of the Barnes maze apparatus under an opaque cylinder. Once the cylinder was removed, rats were left to freely explore the maze for up to 5 minutes or until the target hole was found and entered the escape box. The rats were left in the escape box for 1 minute before being returned to the home cage. In the event the rat did not find the escape hole by the 5-minute mark, it was guided to the hole with a clear beaker and allowed to stay in the beaker, over the hole, for up to 2 minutes. If the rat still did not find the hole in the allotted time, it was gently nudged with the beaker for up to 1 minute. If the rat still did not enter the hole, it was placed in the hole. Once in the escape hole, the rat was left to rest in the escape box for 1 minute. After this minute was up, the rat was returned to the home cage, and the table was cleaned with 70% ethanol. The table was rotated a quarter turn clockwise in preparation for testing the next rat. This was repeated once daily for 4 days (Day 1-4). Probe trials were conducted on Day 8 (3 days after the last training trial). During the probe trial, the escape box was removed, and animals freely explored the maze for 2 minutes. The latency to find the target hole and the time spent in the quadrant containing the target hole were measured and analyzed. After the completion of Probe trial 1, the rats were trained to a new target hole with the same procedure, from Day 9 to 12. Probe trial 2 was conducted on Day 16, and the data were analyzed the same as Probe trial 1.

### Immunohistochemical analysis

Immunohistochemistry was performed as previously described [22]. Briefly, rats from low dose/high stress study were anesthetized with a mixture of Xylazine and Ketamine and then perfused with ice cold phosphate buffered saline (PBS; 137 mM NaCl (Fisher Scientific Cat# S271), 2.7 mM potassium chloride (KCl; Fisher Scientific Cat# P217), 10 mM sodium phosphate dibasic (HNa2PO_4_; Fisher Scientific Cat# S393), 1.8 mM monobasic potassium phosphate (KH_2_PO_4_; Fisher Scientific Cat# BP362); pH 7.4) followed by ice cold 4% paraformaldehyde solution (PFA; Fisher Scientific Cat# T353). Brains were dissected out and left to post-fixing in 4% PFA at 4°C for at least one week. After fixing, brains were moved to a cryoprotectant solution consisting of 30% sucrose (PR1MA Cat# KCS24060) and 0.02% sodium azide (NaN_3_; Fisher Scientific Cat# S227) in PBS and left at 4°C for at least three days, or until the brains sunk to the bottom of the solution. Using a sliding microtome (Leica, SM2010R), the coronal sections were sliced at a 30 μm thickness and collected serially (every 10^th^ section) from the start of the hippocampus and going until no more hippocampal sections could be obtained. Sections were stored in PBS with 0.02% NaN_3_ at 4°C. Sections were blocked in 5% normal goat serum (Vector Laboratories Cat# S-1000-20) with 0.2% Triton-X100 (Sigma-Aldrich Cat# 9002-93-1) in PBS. Primary antibodies used for immunohistochemical staining included the following: glial fibrillary acidic protein (GFAP; 1:1000; Sigma-Aldrich Cat# G3893), parvalbumin (PV; 1:1000; Sigma-Aldrich Cat# P3088) and doublecortin (DCX; 1:500; Abcam Cat# ab18723). Secondary antibodies were purchased from Fisher Scientific and included Alexa Fluor 594 goat anti-mouse (1:200; Invitrogen Cat# A-11032), anti-rabbit IgG (1:200; Invitrogen Cat# A-11037), and DAPI (1:5000; Thermo Fisher Scientific Cat# D1306, RRID: AB_2629482). Images were taken with a fluorescent Axioskop 2 plus microscope (×10/0.25NA Achroplan, Carl Zeiss) using AxioVision (v4.8) software and analyzed with FIJI software [22]. Whole images of DG and HC were stitched together with a stitching plugin found within the FIJI software [27].

PV^+^ cells were counted manually and divided by the area of the entire hippocampus region. Data were averaged across all sections stained for all animals in that group (>5 sections per animal, n=4 animals per group). DCX^+^ cells were counted manually after a brief digital processing on FIJI software. Briefly, images of whole hippocampi were outlined and cropped. Under the analyze tab the scale was set in accordance with our device’s appropriate scale setting (1.526 pixels = 1.00 μm). The areas of the hippocampi were found using the measure tool. After this, the image type was changed to 8-bit, the background was subtracted with a rolling ball radius of 30 and image threshold was adjusted to 0.18%. Images were then analyzed via the “analyze particles” function with the size (pixel^2) setting set to 35-Infinity. Cells found outside of the hippocampus were removed from the final count. Data were averaged across all sections stained for all animals in that group (≥4 sections per animal, n=4 animals per group).

GFAP^+^ cells were analyzed in the dentate gyrus (DG), CA1, and CA3 of the hippocampi using FIJI software (ImageJ) to process the images and to automatically count the astrocytes. For the DG analysis, images were uploaded to FIJI, channels were split, and the red channel was analyzed. The scale was set as previously mentioned. Following this, the image type was changed to 8-bit, and the background was subtracted using a rolling ball radius of 30. The threshold was set to MaxEntropy with “White Objects on Black Background” checked. We analyzed three non-overlapping 200 x 200 pixel squares in the middle of the DG that contained only astrocytes. Particles were then analyzed with the “Analyze Particles” function, with the size set to 35-Infinity pixels and “Exclude on Edges” checked. For the CA1 analysis, after setting the scale, the area was cropped to a 1200 x 500 pixel rectangle. The background was subtracted with a rolling ball radius of 30, and the threshold was set to RenyiEntropy with “White Objects on Black Background” checked. Particles were analyzed using the “Analyze Particles” function with the size set to 5-Infinity pixels and “Exclude on Edges” checked. The CA3 analysis followed a similar procedure to CA1. The area was cropped to a 500 x 900 pixel rectangle, the background was subtracted, and the threshold was set to RenyiEntropy. Particles were then analyzed using the same settings as for CA1.

### Western blot analysis

Hippocampal tissues were placed in 10x volume of homogenization buffer and homogenized in 20 consecutive strokes using a motorized overhead stirrer (SouthwestScience Cat# SOS20) with a 2 ml glass mortar and polytetrafluoroethylene pestle. The homogenization buffer contained 0.32 M Sucrose (PR1MA Cat# KCS24060); 10 mM Tris (pH 7.5; Bio-Rad Cat# 161-0715); 1 mM EDTA (pH 8.0; Fisher Scientific Cat# BP120); 1 mM EGTA (VWR Cat# 0732); and 1x Protease Inhibitor (ThermoFisher Scientific Cat# A32965). Total cell lysate was sonicated (Fisher Scientific Cat# F60 Sonic Dismembrator) prior to reading the protein concentration using a DC™ Protein Assay Kit I (Bio-Rad Cat# 5000111). Samples were prepared and run on 8% SDS-PAGE polyacrylamide gels. Proteins were transferred from the gels onto nitrocellulose membranes using Bio-Rad’s Trans-Blot Turbo Transfer System (Bio-Rad Cat# 1704150). Membranes were blocked for 30 minutes in phosphate-buffered saline mixed with 0.1% Tween-20 (Fisher Scientific Cat# BP337-100) and 5% non-fat dried milk (Kroger, UPC: 0001111008736). Once blocked, membranes were probed overnight with primary antibody. The following antibodies were used: EAAT2 (1:20,000; custom antibody; [28]); VGLUT1 (1:500; EMD Millipore Corp. Cat# MAB5502); PSD95 (1:1000-2000; Invitrogen Cat# MA1-046); actin (1:10,000; EMD Millipore Corp. Cat# MAB1501); synaptophysin (1:500; Cell Signaling Technology Cat# 9020S); GFAP (1:1000; Millipore Sigma Cat# G3893); NCAM (1:1000; Cell Signaling Technology Cat# 99746S); gephyrin (1:500; GeneTex Cat# GTX109734); VGAT (1:2000; Invitrogen Cat# PA5-96231); and doublecortin (1:500; Abcam Cat# 18723). Membranes were subsequently washed with phosphate buffered saline with 0.1% Tween-20 (PBST) and incubated in their respective secondary antibodies for one hour. Membranes were once again washed in PBST before probing with WesternBright ECL (Thomas Sci Cat# C942A85) or WesternBright Sirius (Thomas Sci Cat# C942A92) according to manufacturer’s instructions. Blots were imaged on the ChemiDoc™ Touch Imaging System (Bio-Rad Cat# 1708370). Band intensities were quantified with Image Lab software (Bio-Rad, V 5.2.1) then normalized to loading controls (GFAP or actin). Results are presented as fold-change relative to controls.

### Statistical analysis

We used GraphPad Prism (v5.01) to perform all data analysis and make all graphs. All n-values are shown in the figure legends. Body weight data were analyzed using two-way ANOVA with a Bonferroni post hoc test. Behavior data were analyzed using both one-way and two-way ANOVA. For one-way ANOVA, a Bartlett’s omnibus test was used to determine if unequal variances were present. If the one-way ANOVA was significant and no unequal variances were found, it was followed by a Tukey’s post hoc test. If significant variances were present, a Kruskal-Wallis test followed by a Dunn’s post hoc test was used. Two-way ANOVA were followed by a Bonferroni post hoc test. Data from non-responders were excluded from the behavioral datasets. Immunohistochemical data were also analyzed using both one-way and two-way ANOVA, as described above. For these analyses, we used hippocampal sections from: 4 animals for DCX immunostaining (≥ 5 sections per animal); 4 animals for PV (≥ 4 sections per animal); and 3 animals for GFAP (≥ 3 sections per animal). Western blot data were analyzed by converting the signal to fold-change relative to the control vehicle. A one-sample t-test was used to compare these values to the theoretical value of 1.0. All data are reported as mean ± standard deviation (SD) with p-values less than 0.05 considered significant.

## Results

We previously evaluated compound LDN/OSU-215111 in a mouse model of GWI and observed significant beneficial effects [22]. After this study, we found that LDN/OSU-215111 is rapidly metabolized into a major metabolite upon entering the body in rodents. Further investigation revealed that this metabolite exhibits good brain penetration, oral bioavailability, pharmacokinetics, potency, solubility, stability, and specificity. We named this metabolite LH-001. To validate our previous findings in mice, this study aimed to investigate the effects of LH-001 in rats exposed to GW-related chemicals and stress. All behavioral raw data, immunohistochemical raw data, and Western blotting raw data can be found in S1, S2, and S3 files, respectively.

We first investigated LH-001 using the “low dose/low stress exposure” protocol (**Table 1**), based on the method reported by Parihar et al. [23]. Male Sprague Dawley rats were randomly divided into four groups (n=12/group): control/vehicle (no GW-exposure, treated with vehicle), control/LH-001 (no GW-exposure, treated with LH-001), GWI/vehicle (GW-exposure, treated with vehicle), and GWI/LH-001 (GW-exposure, treated with LH-001). Rats in the GWI/vehicle and GWI/LH-001 groups were exposed to GW chemical agents and stress every weekday for five weeks (described in detail in the Methods section). Body weight was measured weekly over the first six weeks, and the data were analyzed by two-way ANOVA. Starting 4-5 weeks post-GW exposure, rats received either LH-001 or vehicle (described in detail in the Methods section). Behavioral tests assessing anxiety and cognitive functions were performed starting from 8–9 weeks post-GW exposure. LH-001 or vehicle treatment continued throughout the testing period. Four tests were conducted to evaluate anxiety-like behaviors: open field, light and dark transition, elevated plus maze, and novelty-suppressed feeding tests (Fig 1B-H). We assessed the cognitive state of the rats using novel object location, novel object recognition, and Barnes maze tests (Fig 1I-O). All behavioral data were first analyzed by Bartlett’s test to check if variances were equal across four groups, and the results showed unequal variances for the following behavioral data: open field distance traveled (**Fig 1C**), dark and light transition latency (**Fig 1E**), novelty-suppressed feeding latency (**Fig 1H**), and Barnes maze latency (**Fig 1M**) (*p* < 0.05). These data were analyzed using a Kruskal-Wallis test with Dunn’s post hoc test to determine significance. The rest of the data had equal variances and were analyzed using a one-way ANOVA followed by Tukey’s post hoc tests. A two-way ANOVA was also conducted for all behavioral data.

**Fig 1.**
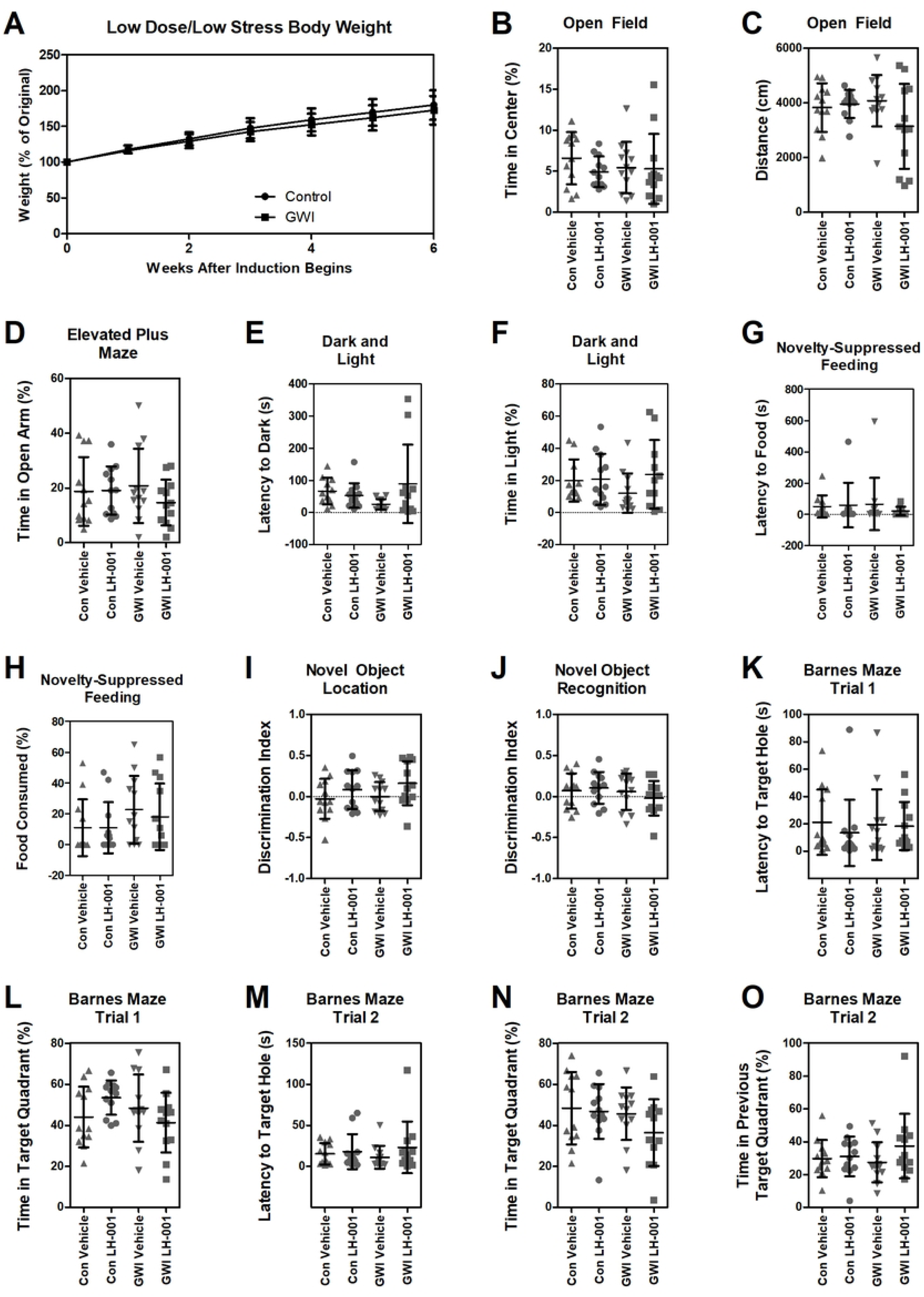
Low dose/low stress exposure does not alter body weight, anxiety, or cognition in rats. Rats were exposed to a low dose of chemical agents and a low-stress protocol for five weeks. Body weight was monitored during and post-exposure. At 4-5 weeks post-exposure, rats received either LH-001 or vehicle. Anxiety-like and cognitive behaviors were assessed starting from 8–9 weeks post-exposure. A two-way ANOVA was used for body weight analysis. Both one-way and two-way ANOVA were used for behavioral analysis. One-way ANOVA analyzed data are shown (B-O). Unless otherwise specified, sample sizes were n = 12 per group. **(A) Body weight.** Body weights were measured weekly over the first six weeks (n = 24/group). No significant differences were found between groups. **(B, C) Open field test.** Time spent in the center (B) and total distance traveled (C) were measured. **(D) Elevated plus maze test.** The percentage of time spent in the open arms was analyzed. **(E-F) Dark and light test.** Latency to enter the dark chamber (E) and percent of time spent in the light chamber (F) were assessed (n = 11–12/group). **(G-H) Novelty-suppressed feeding test.** Latency to begin feeding (G) and amount of food consumed (H) were analyzed (n = 10–12/group). **(I, J) Recognition memory test.** Discrimination index scores for novel object location (I) and novel object recognition (J) were calculated. **(K-L) Barnes maze test.** Latency to find the escape hole (K) and time spent in the target quadrant (L) during the initial training phase were analyzed. **(M-N) Barnes maze re-training.** Latency to the new escape hole (M) and time spent in the new target quadrant (N) were analyzed one week later. **(O) Barnes maze memory recall.** Time spent in the original target quadrant during the re-training phase was analyzed. No significant differences were found between groups for any behavioral test.

No differences in body weight were observed between GW-exposed and control groups at any measured time point (**Fig 1A**). The analyzed anxiety data are shown in Fig 1B-H. In the open-field test, we did not observe a decrease in time spent in the center (**Fig 1B**) or a reduction in total distance traveled (**Fig 1C**), both of which would be expected in an anxiety-like phenotype. This lack of anxiety-like behavior was further supported by the elevated plus maze test, where both the control and GWI groups spent equal amounts of time in the lighted arms (**Fig 1D**). This contrasts with the original model description, which reported GWI rats spending significantly less time in the open arm [29]. Furthermore, GWI rats did not show significantly reduced time to the dark box (**Fig 1E**) or spend less time in the light box (**Fig 1F**) in the dark and light transition test. In the novelty-suppressed feeding test, GWI rats did not approach the novel food more slowly (**Fig 1G**), nor did they consume smaller amounts (**Fig 1H**). A two-way ANOVA showed no main effect of the GW-exposure factor across all anxiety behavioral tests. Collectively, these data demonstrated no evidence of an anxiety-like phenotype.

The analyzed cognition data are shown in Fig 1I-O. GWI rats showed no recognition memory deficits, discriminating effectively between objects in novel and familiar locations (**Fig 1I**) and between novel and familiar objects (**Fig 1J**) 24 hours after initial exposure. In the Barnes maze, GWI rats also exhibited no deficits in spatial memory and learning, with no increase in latency to find the target hole (**Fig 1K**) or decrease in time in the target quadrant (**Fig 1L**) during the test conducted a week after training. Furthermore, GWI rats spent a similar amount of time as controls finding a new trained target hole (**Fig 1M**) and exploring the new (**Fig 1N**) or previous target quadrants (**Fig 1O**). A two-way ANOVA revealed a significant main effect of LH-001 treatment in novel object location data (*F*(1,44) = 4.09, *p* = 0.0493); Fig 1I). This suggests that LH-001 treatment improved the ability to remember and recognize a change in an object’s position, regardless of GW-exposure. Overall, there was no evidence of cognitive deficits in low dose/low stress exposed rats, and LH-001 treatment appeared to improve spatial memory.

Without a clear phenotype in the GWI model, it is impossible to assess whether LH-001 has the potential to provide efficacy for GWI. Therefore, we significantly increased the dose of PB (13 mg/kg), PER (2.6 mg/kg), and DEET (215 mg/kg) [24–25] while maintaining the same five-minute restraint stress (the “high dose/low stress exposure” protocol in Table 1). We performed the same procedures and behavioral assessments. Two-way ANOVA of body weight data showed no differences in body weight between GW-exposed and control groups at any measured time point (**Fig 2A**). Bartlett’s test revealed unequal variances for the following behavioral data: novelty-suppressed feeding latency (**Fig 2G**), Barnes maze trial 1 quadrant (**Fig 2L**) and trial 2 latency (**Fig 2M**). These data were analyzed using a Kruskal-Wallis test with Dunn’s post hoc test. The remaining data were analyzed using a one-way ANOVA with Tukey’s post hoc tests. The analyzed anxiety data (**Fig 2B–H**) showed no indication that increased chemical doses induced anxiety-like phenotypes in the open field task (**Fig 2B-C**), the elevated plus maze (**Fig 2D**), the dark and light test (**Fig 2E-F**), or the novelty suppressed feeding test (**Fig 2G-H**). Interestingly, GWI groups appeared to exhibit less anxiety-like phenotypes than controls. Specifically, GWI/vehicle rats spent significantly more time in the light area of the dark and light test (**Fig 2F**), while GWI/LH-001 rats ate significantly more novel food in the novelty-suppressed feeding test (**Fig 2H**) (*p* < 0.05). In each respective test, the other GWI group (GWI/LH-001 in the dark/light test and GWI/vehicle in the novelty-suppressed feeding test) also showed a tendency toward less anxiety-like behavior, although this did not reach significance. Furthermore, the cognitive tests (**Fig 2I–O**) revealed no cognitive issues in the novel object location (**Fig 2I**) and recognition tasks (**Fig 2J**), or the Barnes maze (**Fig 2K-O**). Overall, increased doses of the chemical agents did not result in an anxiety-like phenotype or cognitive deficits.

**Fig 2.**
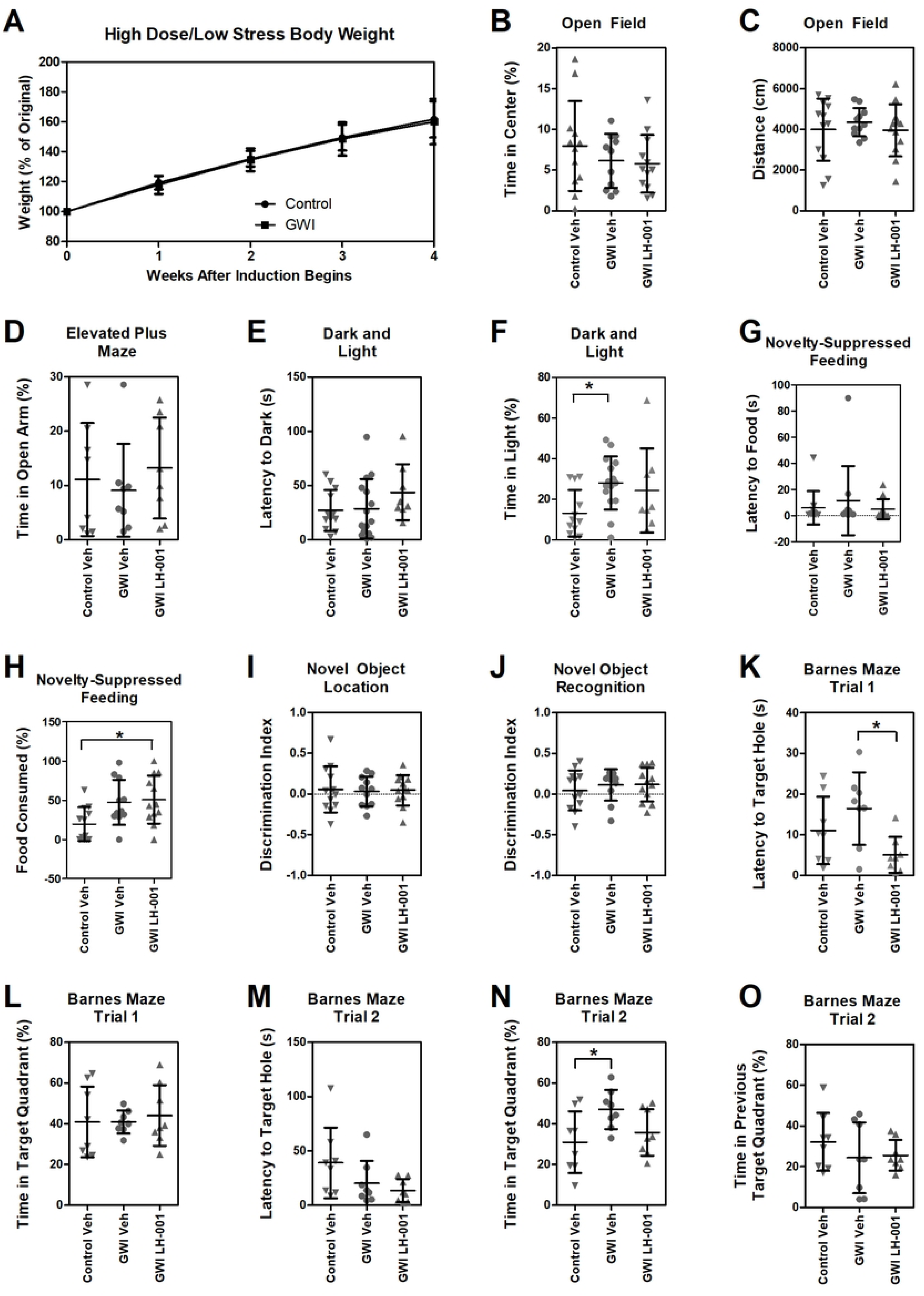
High dose/low stress exposure does not alter body weight or cognition but appears to reduce anxiety in rats. Rats were exposed to a high dose of chemical agents and a low-stress protocol for five weeks. Body weight was monitored during GW exposure. At 4-5 weeks post-exposure, rats received either LH-001 or vehicle. Anxiety-like and cognitive behaviors were assessed starting from 8–9 weeks post-exposure. A two-way ANOVA was used for body weight analysis. A one-way ANOVA was used for behavioral analysis. **(A) Body weight.** Body weights were monitored weekly over the first four weeks (n = 8 for control and n = 16 for GWI). No significant differences were found between groups. **(B, C) Open field test.** Time spent in the center (B) and total distance traveled (C) were measured (n = 12/group). No significant differences were found between groups. **(D) Elevated plus maze test.** The percentage of time spent in the open arms was analyzed (n = 8/group). No significant differences were found between groups. **(E-F) Dark and light test.** Latency to enter the dark chamber (E) and percent of time spent in the light chamber (F) were assessed (n = 8-15/group). GWI/vehicle rats spent significantly more time in the light area; GWI/LH-001 rats also spent more time in the light area but did not reach significance. **(G-H) Novelty-suppressed feeding test.** Latency to begin feeding (G) and amount of food consumed (H) were analyzed (n = 11-12/group). GWI/LH-001 rats ate significantly more food than control/vehicle rats; GWI/vehicle rats also ate more food but did not reach significance. Overall, high dose/low stress exposure appeared to reduce anxiety. **(I, J) Recognition memory test.** Discrimination index scores for novel object location (I) and novel object recognition (J) were calculated (n = 8/group). No significant differences were found between groups. **(K-L) Barnes maze test.** Latency to find the escape hole (K) and time spent in the target quadrant (L) during the initial training phase were analyzed (n = 8/group). GWI/LH-001 rats spent less time finding the target hole than GWI/vehicle rats. **(M-N) Barnes maze re-training.** Latency to the new escape hole (M) and time spent in the new target quadrant (N) were analyzed one week later (n = 8/group). No significant differences were found between groups. **(O) Barnes maze memory recall.** Time spent in the original target quadrant during the re-training phase was analyzed (n = 8/group). GWI/rats spent significantly more time in the target quadrant than control/vehicle rats. Overall, high dose/low stress exposure did not cause cognitive issues. \**p* < 0.05

Several reports from the Committee on Gulf War and Health indicate that GW veterans experienced multiple war-related psychological stresses, such as deployment and combat stress, and environmental stresses, such as dust storms, intense heat during the day and cold at night, and poor living condition [2]. Stress may potentiate the effects of GW chemical agents in the brain and other systems/organs. Higher levels of stress may produce more obvious GWI-like phenotypes [30–31]. We therefore decided to conduct a study using the protocol of GW chemical agents combined with multiple unpredictable stressors (the “low dose/high stress exposure” protocol in Table 1), similar to the mouse model that we previously established [22].

We measured body weight and performed the same behavioral assessments as above. Two-way ANOVA of body weight data showed a significant reduction in the GW-exposed group’s body weight during the GW exposure period at weeks 2, 4, and 5 (**Fig 3A**). We also observed weight loss in our previous study in mice [22]. Analysis of the behavioral data showed unequal variances (Bartlett’s test, *p* < 0.05) for the following data: open field (time; **Fig 3B**), dark and light (latency and time; **Fig 3E, F**), novelty-suppressed feeding (latency; **Fig 3G**), and Barnes maze (trial 1 latency and trial 2 latency; **Fig 3K, M**). These data were analyzed using a Kruskal-Wallis test with Dunn’s post hoc test. The remaining data were analyzed using a one-way ANOVA with Tukey’s post hoc tests. The analyzed anxiety data are shown in Fig 3B-H. There were no significant differences between the groups for open field (**Fig 3B, C**) and novelty-suppressed feeding (**Fig 3G, H**). However, GWI/LH-001 rats spent significantly more time on the open arm of the elevated plus maze (**Fig 3D**) and in the light box of the dark and light (**Fig 3F**) when compared with control/LH-001 rats. A two-way ANOVA revealed a significant interaction effect (*F* (1, 44) = 6.54, *P* = 0.0141) and a significant model effect (*F* (1, 44) = 6.63, *P* = 0.0135) for the elevated plus maze. This indicates that the LH-001 treatment effect only occurred in GW-exposed rats, not in non-exposed rats. This observation has several possible explanations. (1) The low dose/high stress exposure may sensitize certain neural circuits, like the hypothalamic-pituitary-adrenal axis or specific neurotransmitter systems (e.g., the glutamatergic system). LH-001 may only be effective at reversing the overactive signaling in these sensitized circuits. (2) LH-001 may not be active on its own but requires a specific signaling environment created by the high stress to exert its anxiolytic effect. (3) LH-001 and low dose/high stress exposure may target the same biological system with opposing effects. In this case, the effect of LH-001 could be small, and the elevated stress might put the rats above a behavioral or neurochemical threshold where the treatment’s impact becomes detectable. Taken together, the data indicated no evidence of anxiety-like phenotypes with increasing stress. It is notable that in our previous study, mice that received the same GW exposure exhibited clear anxiety-like phenotypes at three months post-exposure, and these phenotypes persisted for at least one year [22].

**Fig 3.**
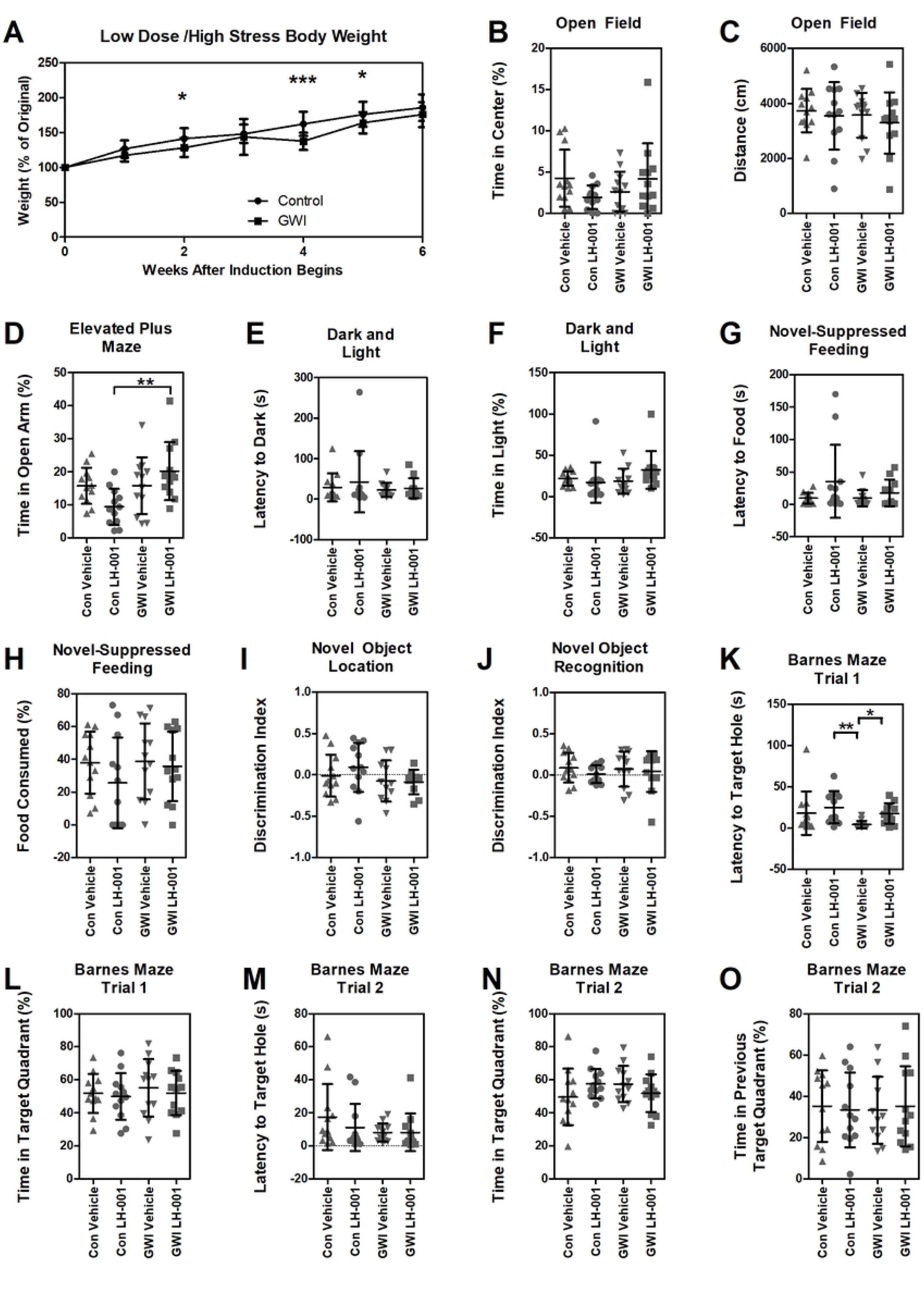
Low dose/high stress exposure reduces body weight but does not cause anxiety or cognitive impairment in rats. Rats were exposed to low doses of chemical agents and a variety of high stressors. Body weight was monitored during and post-GW exposure. At 4 weeks post-exposure, rats received either LH-001 or vehicle. Anxiety-like and cognitive behaviors were assessed starting from 10 weeks post-exposure. A two-way ANOVA was used for body weight analysis. Both one-way and two-way ANOVA were used for behavioral analysis. One-way ANOVA analyzed data are shown (B-O). **(A) Body weight.** Body weights were monitored weekly over the first six weeks (n = 24/group). GW-exposed groups showed a significant reduction in body weight during the GW exposure period. **(B, C) Open field test.** Time spent in the center (B) and total distance traveled (C) were measured (n = 12/group). No significant differences were found between groups. **(D) Elevated plus maze test.** The percentage of time spent in the open arms was analyzed (n = 8/group). GWI/LH-001 rats spent significantly more time in the open arm when compared to control/LH-001 rats. **(E-F) Dark and light test.** Latency to enter the dark chamber (E) and percent of time spent in the light chamber (F) were assessed (n = 8-15/group). No significant differences were found between groups. **(G-H) Novelty-suppressed feeding test.** Amount of food consumed (G) and latency to begin feeding (H) were analyzed (n = 11-12/group). No significant differences were found between groups. **(I, J) Recognition memory test.** Discrimination index scores for novel object location (I) and novel object recognition (J) were calculated (n = 8/group). No significant differences were found between groups. **(K-L) Barnes maze test.** Latency to find the escape hole (K) and time spent in the target quadrant (L) during the initial training phase were analyzed (n = 8/group). GWI/vehicle rats found the target hole more quickly when compared to control/LH-00 rats. **(M-N) Barnes maze re-training.** Latency to the new escape hole (M) and time spent in the new target quadrant (N) were analyzed one week later (n = 8/group). No significant differences were found between groups. **(O) Barnes maze memory recall.** Time spent in the original target quadrant during the re-training phase was analyzed (n = 8/group). Overall, no general trend for anxiety-like phenotypes or cognitive deficits with increasing stress was found. \**p* < 0.05, \*\**p* < 0.01, \*\*\**p* < 0.001

The cognition data are shown in Fig 3I-O. There were no significant differences between the groups for novel object location (**Fig 3I**) and novel object recognition (**Fig 3J**), indicating no spatial memory change. In the Barnes maze (**Fig 3K-O**), GWI/vehicle rats more quickly found the target hole in the initial trial than other groups (**Fig 3K**). A two-way ANOVA revealed a significant main effect for the GWI model on Barnes maze trial 1 latency (K) (*F* (1, 44) = 4.35, *P* = 0.0428). This indicates that low dose/high stress exposed rats found the target hole significantly faster than control rats. The potential reasons for this unexpected result are twofold. First, the low dose/high stress exposure may heighten exploratory drive, causing the GWI rats to search more aggressively and move faster than control rats. Second, it is possible that the low dose/high stress exposure may specifically alter rats’ spatial and learning memory function in a way that benefits this particular task, perhaps by shifting how they prioritize spatial cues. Collectively, the data revealed no evidence of cognitive deficits in rats subjected to the low dose/high stress protocol. Notably, in our previous study, mice under the same protocol exhibited significant cognitive impairments assessed by the same tests [22].

After completing the behavioral assessments, the rats were euthanized for further pathological studies. We focused on the hippocampal regions of the high stress/low dose cohort rats. In our previous study in mice [22], by immunohistochemical staining, we did not observe significant changes for hippocampal pyramidal neurons and microglia regarding the number, morphology, and signal intensity between GW-exposed and control mice. However, we observed significant changes for the following cell types in GW-exposed mice: (i) astrocytes with a reduced area of glial fibrillary acidic protein-positive (GFAP^+^) elements, suggesting astrocytic atrophy; (ii) parvalbumin-positive (PV^+^) interneurons with a reduced number, suggesting their loss; and (iii) doublecortin-positive (DCX^+^) newborn neurons with a reduced number, indicating decreased neurogenesis. We therefore conducted immunohistochemical staining using GFAP, PV, and DCX antibodies to examine astrocytes, PV interneurons, and newborn neurons, respectively. Hippocampal sections from 3–4 animals, with at least 3 sections per animal, were analyzed for each staining. Bartlett’s test revealed unequal variances for the DCX data (*p* < 0.05); these data were analyzed using a Kruskal-Wallis test with Dunn’s post hoc test. The rest of the data were analyzed using a one-way ANOVA with Tukey’s post hoc tests. As shown in Fig 4, the number of GFAP^+^ astrocytes in DG, CA1, and CA3 regions (**Fig 4A–C**) and the number of PV^+^ interneurons in the hippocampus (**Fig 4D**) did not differ significantly between the four groups. The number of DCX^+^ neurons in DG region was significantly reduced in GWI/LH-001 group when compared to control/LH-001 group (**Fig 4E**). A two-way ANOVA revealed a significant main effect of the GWI model for GFAP -CA1 (*F*(1,128) = 4.88, *p* = 0.0289), GFAP -CA3 (*F*(1,131) = 4.88, *p* = 0.0289), and DCX - DG (*F*(1,125) = 11.28, *p* = 0.0010). No significant treatment or interaction effects were found. These results indicated that low dose/high stress exposure significantly reduced hippocampal neurogenesis and that LH-001 treatment had no effect on this reduction. In addition, low dose/high stress exposure altered the number of GFAP^+^ astrocytes in the CA1 and CA3 regions and that LH-001 treatment had no effect on this alteration.

**Fig 4.**
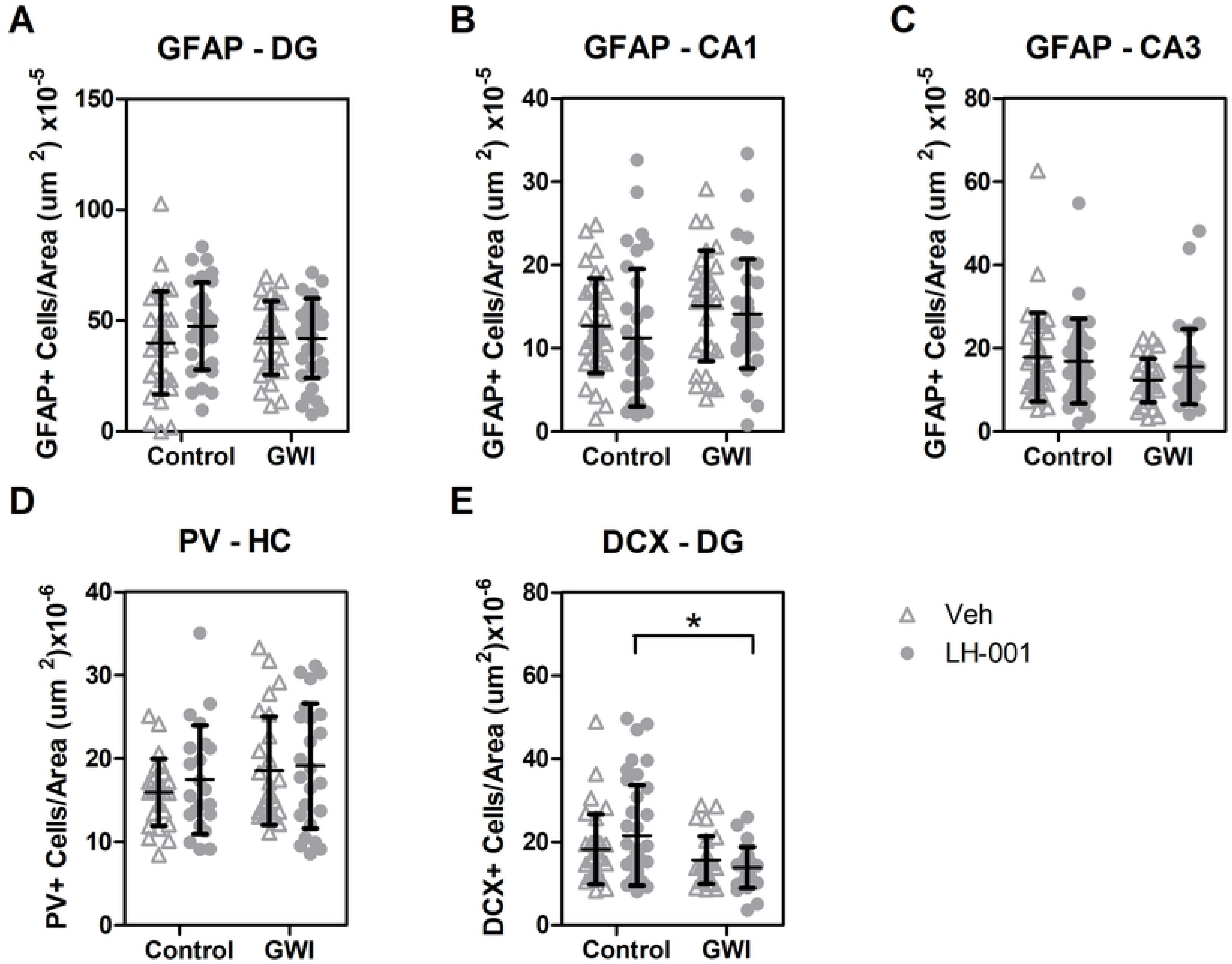
Low dose/high stress exposure reduces hippocampal neurogenesis and does not alter the number of GFAP^+^ astrocyte or PV^+^ interneuron population in rats. Following behavioral assessments (the above low dose/high stress exposure study), a subset of rats was euthanized to examine markers for astrocytes (GFAP^+^), parvalbumin-positive interneurons (PV^+^), and newborn neurons (DCX^+^) in hippocampal regions using immunohistochemical staining. For each staining, 3–4 animals and at least 3 sections per animal were analyzed. Data were analyzed by one-way and two-way ANOVA. One-way ANOVA analyzed data are shown. **(A-C) the number of GFAP^+^ astrocytes**. No significant differences were found between groups (DG, n = 27-33; CA1, n = 31-34; CA3, n = 31-36). **(D) the number of PV^+^ neurons**. No significant differences were found between groups (n = 22-26). **(E) the number of DCX^+^ newborn neurons**. Newborn neurons in DG region were significantly reduced in GWI/LH-001 group compared to the control/LH-001 group (n = 29-39). \**p* < 0.05

We further examined total expression levels of several proteins in hippocampi by Western blot analysis. The proteins studied were: (1) synaptophysin (SYP) and vesicular glutamate transporter 1 (VGLUT1), which are expressed in the presynaptic terminals of glutamatergic neurons; (2) postsynaptic density protein 95 (PSD95) and neural cell adhesin molecule (NCAM), which are expressed in the postsynaptic terminals of glutamatergic neurons; (3) glutamate transporter EAAT2 and GFAP, which are primarily expressed in astrocytes; (4) vesicular GABA transporter (VGAT), which is expressed in the presynaptic terminals of GABAergic neurons; (5) gephyrin (GPHN), which is primarily expressed in the postsynaptic terminals of GABAergic neurons; and (6) doublecortin (DCX), which is expressed in newborn neurons. Results showed that total expression levels of synaptic proteins, including SYP, VGLUT1, PSD95, NCAM, and VGAT, and astrocytic protein, EAAT2, were significantly increased in the control/LH-001 group compared to the control/vehicle group, which reflected LH-001’s effects (**Fig 5**). Notably, the GWI/vehicle group also showed significantly increased total expression levels of VGLUT1 and EAAT2 compared to the control/vehicle group. This suggests that GW exposure may increase glutamate release, and that the observed increase in EAAT2 expression may be a compensatory mechanism to maintain glutamate homeostasis and prevent excitotoxicity. Similarly, VGLUT1 and EAAT2 expression levels were also increased in the GWI/LH-001 group.

**Fig 5.**
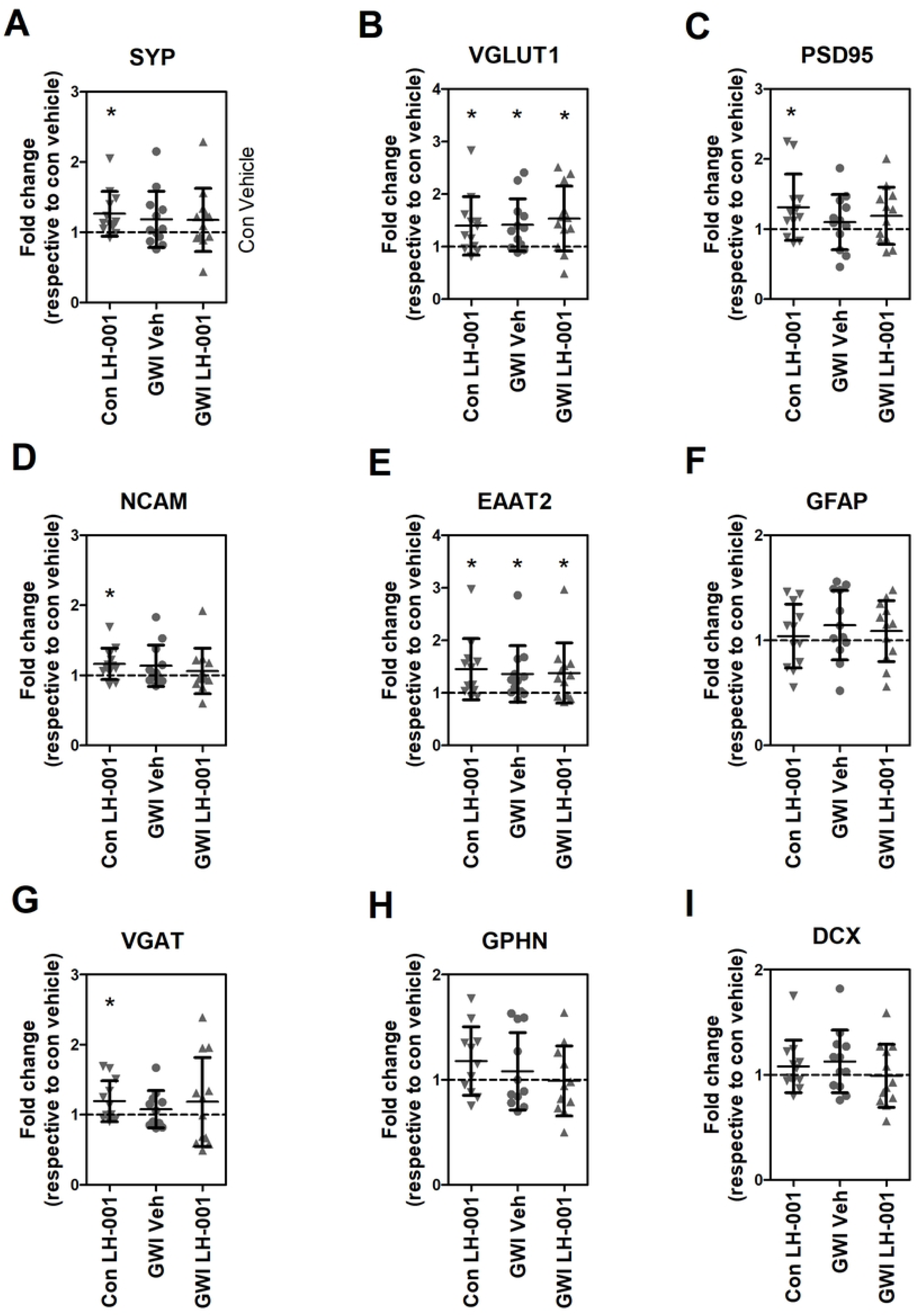
Effects of low dose/high stress exposure and LH-001 treatment on synaptic and astrocytic protein expression in the rat hippocampus. Total cell lysates were prepared from the hippocampus and analyzed for protein expression by Western blotting, followed by quantitative analysis. Results are presented as fold-change relative to the control/vehicle group, which is represented by a dashed line. Data was analyzed by unpaired t-test (n=11-12 each group). The levels of glutamatergic synaptic proteins, including SYP, VGLUT1, PSD95, and NCAM, were significantly increased in the control/LH-001 group compared to the control/vehicle group. The GABAergic synaptic protein VGAT and the astrocytic protein EAAT2 were also significantly increased in the control/LH-001 group compared to the control/vehicle group. Additionally, the levels of VGLUT1 and EAAT2 were significantly increased in the GWI/vehicle and GWI/LH-001 groups compared to the control/vehicle group. Data represents the analysis of six animals per group. Statistical significance was determined using a one-sample t-test (\**p* < 0.05).

## Discussion

This study aimed to test the efficacy of compound LH-001, a novel synapse enhancer, in rat models of GWI. Unexpectedly, our initial GWI induction protocol in rats was ineffective in producing GWI-like phenotypes reported in previous studies [23, 29]. After adjusting the induction protocol twice—once to a more potent chemical exposure and once to a higher level of stress—we were still unable to produce GWI-like phenotypes in rats. These results highlight the challenges of producing a reliable GWI rat model.

These unexpected results prompted a closer examination of existing GWI induction protocols. A comparison of our study’s protocols with published protocols that reported behavioral outcomes is summarized in **Table 3**. The following observations were made. First, the comparison revealed heterogeneity among published protocols that use different combinations of chemical agents, stress, and timelines attempting to produce GWI-like phenotypes. This variability may account for the difficulty researchers face in reproducing similar behavioral outcomes across different laboratories. Even reports from the same laboratory show inconsistent behavioral outcomes. For example, Studies 2 [23] and 3 [29] were reported by the same laboratory using the same protocol. While Study 2 concluded that exposure to GWI-related chemicals and stress did not impair recognition memory, Study 3 found that the same protocol caused recognition memory deficits. Despite using the same protocol, the behavioral outcomes reported in these studies were not replicated in our study (Study 1a). Similarly, Study 7 [32], which also used this protocol, did not observe impaired spatial learning and memory; this was contrary to the results reported in Study 2. In addition, the GWI induction protocols changed even in reports from the same laboratory. For example, Studies 2 [23], 3 [24], 4 [33], and 10 [34] were all reported from the same laboratory. Instead of the 5-min restraint used in Studies 2 and 3, Studies 4 and 10 used a 15-min restraint. Moreover, higher doses of chemical agents were used in Studies 4 and 10. The rationale for these changes is not clearly stated, despite the studies being conducted in the same laboratory. Furthermore, study 12 [35] was able to reproduce similar behavioral outcomes using a dosing and stress regimen similar to that of studies 2 [23] and 3 [24], but the behavioral tests were conducted 1–4-days post-exposure, rather than the typical 3-10 months post-exposure used by other studies. This comparison raises significant concerns regarding the inability to reproduce GWI-like phenotypes in rats and validate the findings across different laboratories. Second, most of the published reports did not clearly indicate the number of animal data included in the statistical analysis. Among the eleven published studies (Study 2-12) listed in Table 3, only Study 6 [36], 10 [34], and 12 [35] showed individual rat data. Study 6 is the only study to present raw behavioral data in the supplemental. Presenting individual data alongside group averages is considered a more thorough and transparent practice in scientific research. It allows other researchers to evaluate the consistency of the findings and observe the full range of responses, including any notable outliers. Third, the central puzzle is our inability to produce GWI-like phenotypes in rats, even with increased doses of chemical agents (Study 1b) or stress (Study 1c). This rules out the possibility that they are the primary reasons. The timing of behavioral assessment is also unlikely to be the reason for the results. In Study 1c (low dose/high stress), we repeated tests at 6–8 months post-exposure, after initially testing at ∼3 months. We still found no GWI-like phenotypes. The strain and source of rats do not appear to be the reason, as different strains—Sprague Dawley or Wistar— from different sources were used across Studies 2-12. There may be other biological or environmental factors underlying the failure to reproduce the GWI-like phenotypes in rats. Taken together, we conclude that GWI-like behavioral phenotypes may not be reliably reproducible in rats.

**Table 3.** GWI Induction Protocols.

| Study | Chemical exposure | Stressor | Testing timing | Stain Sex Source | Conditions | Behavioral Outcomes | Ref |
| --- | --- | --- | --- | --- | --- | --- | --- |
| 1a | PB (1.3 mg/kg)<br>PER (0.13 mg/kg)<br>DEET (40 mg/kg)<br>every weekday for 5 weeks | 5-min restraint every weekday for 5 weeks | Starting from 8-9 weeks post-exposure | Rat/SD male (CRL, USA) | 1. vehicle (no stress)<br>2. chemical + stress (n=12/group) | The chemical + stress group showed no evidence of anxiety-like phenotypes in the **OFT**, **LDT**, **EPMT**, or **NSFT**, nor did it show cognitive deficits in the **OLT**, **NORT**, or **BMT**, compared to the control group.<br>Individual rat data were shown for each behavioral test. | this study |
| 1b | PB (13 mg/kg)<br>PER (2.6 mg/kg)<br>DEET (215 mg/kg) every weekday for 5 weeks | 5-min restraint every weekday for 5 weeks | Starting from 8-9 weeks post-exposure | Rat/SD male (CRL, USA) | 1. vehicle (no stress)<br>2. chemical + stress (n=8-12/group) | The chemical + stress group showed no evidence of anxiety-like phenotypes in the **OFT**, **LDT**, **EPMT**, or **NSFT**, nor did it show cognitive deficits in the **OLT**, **NORT**, or **BMT**, compared to the control group.<br>Individual rat data were shown for each behavioral test. | this study |
| 1c | PB (1.3 mg/kg)<br>PER (0.13 mg/kg)<br>DEET (40 mg/kg)<br>every weekday for 4 weeks | 2 unpredictable stressors every weekday for 4 weeks | Starting from 10 weeks post-exposure | Rat/SD male (CRL, USA) | 1. vehicle (no stress)<br>2. chemical + stress (n=12/group) | The chemical + stress group showed no evidence of anxiety-like phenotypes in the **OFT**, **LDT**, **EPMT**, or **NSFT**, nor did it show cognitive deficits in the **OLT**, **NORT**, or **BMT**, compared to the control group. | this study |
|  |  |  |  |  |  | Individual rat data were shown for each behavioral test. |  |
| 2 | PB (1.3 mg/kg)<br>PER (0.13 mg/kg)<br>DEET (40 mg/kg)<br>daily for 4 weeks | 5-min restraint<br>daily for 4 weeks | Starting<br>from 3<br>months<br>post-exposure | Rat/SD,<br>No gender<br>info<br>provided<br>(Harlan,<br>USA) | 1. vehicle (no stress)<br>2. chemical (no stress)<br>3. chemical + stress<br>(n=6/group) | The chemical (no stress) and chemical + stress groups showed (i) increased immobility time in **FST**, (ii) fewer open-arm entries and reduced open-arm dwell time in **EPMT**, (iii) impaired spatial learning in **MWM**, and (iv) normal recognition memory in **NORT**, compared to the control group.<br>No individual rat data was shown for each behavioral test. | [23] |
| 3 | PB (1.3 mg/kg)<br>PER (0.13 mg/kg)<br>DEET (40 mg/kg)<br>daily for 4 weeks | 5-min restraint<br>daily for 4 weeks | Starting<br>from 3<br>months<br>post-exposure | Rat/SD,<br>No gender<br>info<br>provided<br>(Harlan,<br>USA) | 1. naive control (no exposure)<br>2. chemical + stress<br>(n=16-28/group) | The chemical + stress group exhibited (i) no anxiety-like behavior in **OFT**, (ii) impaired location memory in **OLT**, (iii) impaired recognition memory in **NORT**, (iv) decreased motivation for running on wheel, and (v) longer latency to eat food in **NSFT**, compared to the control group.<br>No individual rat data was shown in each behavioral test. | [29] |
| 4 | PB (2 mg/kg)<br>PER (0.2 mg/kg)<br>DEET (60 mg/kg)<br>daily for 4 weeks | 15-min restraint<br>daily for 4 weeks | Starting<br>from 10<br>months<br>post-exposure | Rat/SD<br>male<br>(Harlan,<br>USA) | 1. naive control (no exposure; n=23 for **NORT**, n=10 for **OIPT**)<br>2. chemical + stress<br>(n=19 for **NORT**, n=16 for **OIPT**) | The chemical + stress group exhibited (i) impaired recognition memory in **NORT** and (ii) impaired object-in-place associative recognition memory in **OIPT**.<br>No individual rat data was shown in each behavioral test. | [33] |
| 5 | PB (1.3 mg/kg)<br>daily for 14 days | 6-hour restraint<br>daily for 10 days | Starting<br>from 12<br>weeks<br>post-exposure | Rat/SD<br>male<br>(Harlan,<br>USA) | 1. vehicle (no stress)<br>2. PB (no stress)<br>3. vehicle + stress<br>4. PB + stress<br>(n=12-15/group) | No significant difference in the time spent exploring the novel object (**NORT**) was found between groups.<br>When challenged with acute LPS, PB-treated rats, irrespective of stress history, spent significantly less time with the novel object in **NORT** and exhibited reduced time in the target quadrant in **MWM**.<br>No individual rat data was shown in each behavioral test. | [37] |
| 6 | PB (1.3 mg/kg)<br>daily and<br>intranasal LPS (50<br>µl of 10 mg/5 ml) | 1 unpredictable<br>stressor daily<br>for 33 days | Starting<br>on day 35<br>post-exposure | Rat/Wistar<br>male &<br>female | 1. naive control<br>(n=14; all males)<br>2. chemical + stress | The chemical + stress group rats spent significantly less time with the novel object in **NORT**, in the open arm in **EPMT**, and | [36] |
|  | every weekday for 33 days |  |  | (Harlan, USA) | (n=36; mixed males & females) | in the center of the open field in **OFT**, compared to the no exposure group. There was no significant difference in the time spent in TST and FST between the groups.<br>• Individual rat data were shown in each behavioral test. |  |
| 7 | PB (1.3 mg/kg)<br>PER (0.13 mg/kg)<br>DEET (40 mg/kg)<br>daily for 4 weeks | 5-min restraint daily for 4 weeks | Starting from 3 months post-exposure | Rat/SD male (CRL, USA) | 1. vehicle (no stress)<br>2. chemical (no stress)<br>3. chemical + stress (n=9-10/group) | • The chemical and chemical + stress groups showed no significant differences in (i) motor response to a sudden, intense sound in **ASR** or (ii) spatial learning and memory in **MWM**, but did show an increase in (iii) motor activity level in **OFT** when compared to the control group<br>• No individual rat data was shown in each behavioral test. | [32] |
| 8 | PB (1.3 mg/kg)<br>daily for 2 weeks | 6-hour restraint daily for 10 days | 10 days post-exposure | Rat/SD male (animal source not provided) | 1. vehicle (no stress)<br>2. vehicle + stress<br>3. PB (no stress)<br>4. PB + stress (n=13-14/group) | • The PB + stress and vehicle + stress groups exhibited deficits in ability to learn and remember an association between a tone and a foot shock in **CFC** test, but the PB alone group did not.<br>• No individual rat data was shown in each behavioral test. | [38] |
| 9 | PB (1.5 mg/kg)<br>every weekday for 2 weeks | **PCA** stress for 25 min every weekday for 2 weeks | Day 52, 113, & 199 post-exposure | Rat/Wistar male (Harlan, France) | 1. vehicle (no stress)<br>2. vehicle + stress<br>3. PB (no stress)<br>4. PB + stress (n=18-20/group) | • The PB + stress group exhibited a learning deficit in **MWM**, which was not observed in the PB or stress group. The PB + stress group also showed significantly increased aggression in **SEA** on day 52 post-exposure, but not on day 113 or 199.<br>• No individual rat data was shown in each behavioral test. | [39] |
| 10 | PB (2.0 mg/kg)<br>PER (0.2 mg/kg)<br>DEET (60 mg/kg)<br>daily for 4 weeks | 15-min restraint daily for 4 weeks | Starting from 8.5 months post-exposure | Rat/SD male (Harlan, USA) | 1. naive control<br>2. chemical + stress (n=10-15/group) | • The chemical + stress group exhibited (i) impaired location memory in **OLT**, (ii) impaired pattern separation memory in **PST**, (iii) anhedonia-like behavior in **SPT**, and (iv) hyperalgesia-like behavior in **VFT**, compared to the control group.<br>• Individual rat data were shown in each behavioral test. | [34] |
| 11 | PB (1.3 mg/kg)<br>PER (0.13 mg/kg)<br>DEET (40 mg/kg)<br>daily for 4 weeks | 5-min restraint<br>daily for 4<br>weeks | 5 and 10<br>months<br>post-exposure | Rat/SD<br>male<br>(Taconic<br>Farms,<br>USA) | 1. naive control<br>(n=10)<br>2. chemical + stress<br>(n=20) | • The chemical + stress group exhibited (i) increased in aggressive behavior, (ii) increased anxiety-like behavior in **OFT** and **EPMT**, (iii) dysfunctional novel object recognition memory in **NORT**, (iv) dysfunctional spatial learning and memory in **MWM**, compared to the control group.<br>• No individual rat data was shown in each behavioral test. | [40] |
| 12 | PB (1.3 mg/kg)<br>PER (0.13 mg/kg)<br>DEET (40 mg/kg)<br>daily for 4 weeks | 5-min restraint<br>daily for 4<br>weeks | Starting<br>from 1-4<br>days post<br>exposure | Rat/SD<br>male<br>(animal<br>source not<br>provided) | 1. control<br>2. chemical + stress<br>3. chemical + stress<br>+ minocycline<br>(n=18/group) | • The chemical + stress group exhibited (i) increased anxiety like behavior in the **OFT** and **EPMT** compared to control and minocycline groups, (ii) memory deficits compared to the other two groups in the **NORT**, and (iii) greater depressive-like behavior in the **FST** compared to the other two groups.<br>• Individual data was presented in each behavioral test. | [35] |

CRL: Charles River Laboratories; LPS: lipopolysaccharide; SD: Sprague Dawley
**ASR**: acoustic startle response; **BMT**: Barnes maze test; **CFC**: cued fear conditioning; **EPMT**: elevated plus maze test; **FST**: forced swim test; **LDT**: light and dark transition; **MWM**: Morris water maze; **NORT**: novel object recognition test; **NSFT**: novelty-suppressed feeding test; **OFT**: open field test; **OIPT**: object-in-place associative recognition memory test; **OLT**: object location test; **PCA**: pole climbing avoidance; **PST**: pattern separation test; **SEA**: shock-elicited aggression test; **SPT**: Sucrose preference test; **TST**: tail suspension test; **VFT**: Von Frey test

*In-vivo* testing in animal models is a regulatory requirement and an essential preclinical step before human clinical trials. The minimal requirement for a valid animal model is that it must reliably reproduce one or more aspects of a human disease or biological process. However, published animal studies often lack reproducibility due to several potential reasons, including inherent biological variability in animals and their environments, lack of transparent reporting, flaws in data analysis and statistics, publication bias favoring positive results, and researcher incentives that favor novelty over rigor. Enhancing the rigor and reproducibility of animal studies is paramount for the GWI scientific community.

## Supporting Information

**S1 File. Behavioral raw data.** Behavioral raw data for all groups of rats for all behavioral tests including, open field, dark and light, elevated plus maze, novelty-suppressed feeding, novel object location, novel object recognition, and Barnes maze.

**S2 File. Immunohistochemical raw data.** Immunohistochemical raw data and representative images for the number of antibody positive cells per area for glial fibrillary acidic protein, doublecortin, and parvalbumin.

**S3 File. Western blotting raw data.** Western blotting raw data with images. Western blots were analyzed by choosing an exposure time that was below saturation for each respective protein.

## References

1. GWIRP. The Gulf War Illness Landscape. Department of Defense; 2020. Available from: https://cdmrp.health.mil/gwirp/pdfs/GWIRP_Landscape_2020.pdf

2. Cory-Slechta D, Wedge R. Gulf War and Health: Volume 10: Update of Health Effects of Serving in the Gulf War, 2016. Washington, D.C.: National Academies Press. 2016: 21840. doi:10.17226/21840

3. Gray GC. Self-reported Symptoms and Medical Conditions among 11,868 Gulf War-era Veterans: The Seabee Health Study. Am J Epidemiol. 2002;155: 1033–1044. doi:10.1093/aje/155.11.1033

4. Kang HK, Li B, Mahan CM, Eisen SA, Engel CC. Health of US Veterans of 1991 Gulf War: A Follow-Up Survey in 10 Years. J Occup Environ Med. 2009;51: 401–410. doi:10.1097/JOM.0b013e3181a2feeb

5. Toomey R, Alpern R, Vasterling JJ, Baker DG, Reda DJ, Lyons MJ, et al. Neuropsychological functioning of U.S. Gulf War veterans 10 years after the war. J Int Neuropsychol Soc. 2009;15: 717–729. doi:10.1017/S1355617709990294

6. White RF, Steele L, O’Callaghan JP, Sullivan K, Binns JH, Golomb BA, et al. Recent research on Gulf War illness and other health problems in veterans of the 1991 Gulf War: Effects of toxicant exposures during deployment. Cortex. 2016;74: 449–475. doi:10.1016/j.cortex.2015.08.022

7. Fulco CE, Liverman CT, Sox HC. Gulf War and Health: Volume 1. Depleted Uranium, Pyridostigmine Bromide, Sarin, and Vaccines. Washington, D.C.: National Academies Press; 2000: 9953. doi:10.17226/9953

8. Haley RW, Kurt TL. Self-reported exposure to neurotoxic chemical combinations in the Gulf War. A cross-sectional epidemiologic study. JAMA. 1997;277(3): 231–237. doi:10.1001/jama.1997.03540270057027

9. Steele L, Sastre A, Gerkovich MM, Cook MR. Complex factors in the etiology of Gulf War illness: wartime exposures and risk factors in veteran subgroups. Environ Health Perspect. 2012;120: 112–118. doi:10.1289/ehp.1003399

10. Wolfe J, Proctor SP, Erickson DJ, Hu H. Risk factors for multisymptom illness in US Army veterans of the Gulf War. J Occup Environ Med. 2002;44: 271–281. doi:10.1097/00043764-200203000-00015

11. Wang X, Ali N, Lin CG. Emerging role of glutamate in the pathophysiology and therapeutics of Gulf War illness. Life Sci. 2021;280: 119609. doi:10.1016/j.lfs.2021.119609

12. Golomb BA. Acetylcholinesterase inhibitors and Gulf War illnesses. Proceedings of the National Academy of Sciences of the United States of America. 2008; 105: 4295–4300. doi:10.1073/pnas.0711986105

13. Meyer, DA, Shafer, TJ. Permethrin, but not deltamethrin, increases spontaneous glutamate release from hippocampal neurons in culture. Neurotoxicology. 2006;27: 594–603. doi:10.1016/j.neuro.2006.03.016

14. Pavlovsky L, Browne, RO, Friedman A. Pyridostigmine enhances glutamatergic transmission in hippocampal CA1 neurons. Experimental neurology. 2003;179: 181–187. doi:10.1016/s0014-4886(02)00016-x

15. Jaiswal C, Singh AK. Particulate matter exposure and its consequences on hippocampal neurogenesis and cognitive function in experimental models. Environ Pollut. 2024 Dec 15;363(2):125275. doi:10.1016/j.envpol.2024.125275.

16. Brooks AW, Sandri BJ, Nixon JP, Nurkiewicz TR, Barach P, Trembley JH, Butterick TA. Neuroinflammation and Brain Health Risks in Veterans Exposed to Burn Pit Toxins. Int J Mol Sci. 2024 Sep 10;25(18):9759. doi:10.3390/ijms25189759.

17. Vellingiri B. A deeper understanding about the role of uranium toxicity in neurodegeneration. Environ Res. 2023 Sep 15;233: 116430. doi:10.1016/j.envres.2023.116430.

18. Tomasini MC, Beggiato S, Ferraro L, Tanganelli S, Marani L, Lorenzini L, Antonelli T. Prenatal exposure to 2,3,7,8-tetrachlorodibenzo-p-dioxin produces alterations in cortical neuron development and a long-term dysfunction of glutamate transmission in rat cerebral cortex. Neurochem Int. 2012 Oct;61(5): 759–66. doi:10.1016/j.neuint.2012.07.004.

19. de la Monte SM, Tong M. Agent Orange Herbicidal Toxin-Initiation of Alzheimer-Type Neurodegeneration. J Alzheimers Dis. 2024;97(4): 1703–1726. doi: 10.3233/JAD-230881.

20. Kassem MS, et al. Stress-induced grey matter loss determined by MRI is primarily due to loss of dendrites and their synapses. Molecular neurobiology 2013;47: 645:661 doi:10.1007/s12035-012-8365-7

21. Treccani, G, et al. Stress and corticosterone increase the readily releasable pool of glutamate vesicles in synaptic terminals of prefrontal and frontal cortex. Molecular psychiatry.2014;19: 433–443. doi:10.1038/mp.2014.5

22. Wang X, Xu Z, Zhao F, Lin KJ, Foster JB, Xiao T, et al. Restoring tripartite glutamatergic synapses: A potential therapy for mood and cognitive deficits in Gulf War illness. Neurobiol Stress. 2020;13: 100240. doi:10.1016/j.ynstr.2020.100240

23. Parihar VK, Hattiangady B, Shuai B, Shetty AK. Mood and memory deficits in a model of Gulf War illness are linked with reduced neurogenesis, partial neuron loss, and mild inflammation in the hippocampus. Neuropsychopharmacology. 2013;38(12): 2348–2362. doi:10.1038/npp.2013.158

24. Abou-Donia M. Co-exposure to pyridostigmine bromide, DEET, and/or permethrin causes sensorimotor deficit and alterations in brain acetylcholinesterase activity. Pharmacol Biochem Behav. 2004;77: 253–262. doi:10.1016/j.pbb.2003.10.018

25. Flunker LK, Nutter TJ, Johnson RD, Cooper BY. DEET potentiates the development and persistence of anticholinesterase dependent chronic pain signs in a rat model of Gulf War Illness pain. Toxicol Appl Pharmacol. 2017;316: 48–62. doi:10.1016/j.taap.2016.12.014

26. Xu Z, Foster JB, Lashley R, Wang X, Benson E, Kidd G, et al. Impact of a pyridazine derivative on tripartite synapse ultrastructure in hippocampus: a three-dimensional analysis. Front Cell Neurosci. 2023;17: 1229731. doi:10.3389/fncel.2023.1229731

27. Preibisch S, Saalfeld S, Tomancak P. Globally optimal stitching of tiled 3D microscopic image acquisitions. Bioinformatics. 2009;25: 1463–1465. doi:10.1093/bioinformatics/btp184

28. Guo H, Lai L, Butchbach ME, Stockinger MP, Shan X, Bishop GA, Lin CL. Increased expression of the glial glutamate transporter EAAT2 modulates excitotoxicity and delays the onset but not the outcome of ALS in mice. Hum Mol Genet. 2003;12(19):2519–32. doi: 10.1093/hmg/ddg267

29. Hattiangady B, Mishra V, Kodali M, Shuai B, Rao X, Shetty AK. Object location and object recognition memory impairments, motivation deficits and depression in a model of Gulf War illness. Front Behav Neurosci. 2014 Mar 13;8: 78. doi:10.3389/fnbeh.2014.00078

30. Abdel-Rahman A, Abou-Donia SM, El-Masry EM, Shetty AK, Abou-Donia MB. Stress and Combined Exposure to Low Doses of Pyridostigmine Bromide, DEET, and Permethrin Produce Neurochemical and Neuropathological Alterations in Cerebral Cortex, Hippocampus, and Cerebellum. J Toxicol Environ Health A. 2004;67: 163–192. doi:10.1080/15287390490264802

31. Macht VA, Woodruff JL, Maissy ES, Grillo CA, Wilson MA, Fadel JR, et al. Pyridostigmine bromide and stress interact to impact immune function, cholinergic neurochemistry and behavior in a rat model of Gulf War Illness. Brain Behav Immun. 2019;80: 384–393. doi:10.1016/j.bbi.2019.04.015

32. Gargas NM, Ethridge VT, Miklasevich MK, Rohan JG. Altered hippocampal function and cytokine levels in a rat model of Gulf War illness. Life Sci. 2021;274: 119333. doi:10.1016/j.lfs.2021.119333

33. Madhu LN, Attaluri S, Kodali M, Shuai B, Upadhya R, Gitai D, et al. Neuroinflammation in Gulf War Illness is linked with HMGB1 and complement activation, which can be discerned from brain-derived extracellular vesicles in the blood. Brain Behav Immun. 2019;81: 430–443. doi:10.1016/j.bbi.2019.06.040

34. Kodali M, Madhu LN, Kolla VSV, Attaluri S, Huard C, Somayaji Y, et al. FDA-approved cannabidiol [Epidiolex®] alleviates Gulf War Illness-linked cognitive and mood dysfunction, hyperalgesia, neuroinflammatory signaling, and declined neurogenesis. Mil Med Res. 2024;11: 61. doi:10.1186/s40779-024-00563-2

35. Cheng C-H, Guan Y, Chiplunkar VP, Mortazavi F, Medalla ML, Sullivan K, et al. Nerve agent exposure and physiological stress alter brain microstructure and immune profiles after inflammatory challenge in a long-term rat model of Gulf War Illness. Brain Behav Immun-Health. 2024;42: 100878. doi:10.1016/j.bbih.2024.100878

36. Keledjian K, Tsymbalyuk O, Semick S, Moyer M, Negoita S, Kim K, et al. The peroxisome proliferator-activated receptor gamma (PPARγ) agonist, rosiglitazone, ameliorates neurofunctional and neuroinflammatory abnormalities in a rat model of Gulf War Illness. PLOS ONE. 2020;15: e0242427. doi:10.1371/journal.pone.0242427

37. Burzynski HE, Ayala KE, Frick MA, Dufala HA, Woodruff JL, Macht VA, et al. Delayed cognitive impairments in a rat model of Gulf War Illness are stimulus-dependent. Brain Behav Immun. 2023;113: 248–258. doi:10.1016/j.bbi.2023.07.003

38. Macht VA, Woodruff JL, Maissy ES, Grillo CA, Wilson MA, Fadel JR, et al. Pyridostigmine bromide and stress interact to impact immune function, cholinergic neurochemistry and behavior in a rat model of Gulf War Illness. Brain Behav Immun. 2019;80: 384–393. doi:10.1016/j.bbi.2019.04.015

39. Lamproglou I, Barbier L, Diserbo M, Fauvelle F, Fauquette W, Amourette C. Repeated stress in combination with pyridostigmine. Behav Brain Res. 2009;197: 301–310. doi:10.1016/j.bbr.2008.08.03130.

40. Wu X, Shetty AK, Reddy DS. Long-term changes in neuroimaging markers, cognitive function and psychiatric symptoms in an experimental model of Gulf War Illness. Life Sci. 2021;285: 119971. doi:10.1016/j.lfs.2021.119971

